# Unexpected diversity in socially synchronized rhythms of shorebirds

**DOI:** 10.1101/084806

**Authors:** Martin Bulla, Mihai Valcu, Adriaan M. Dokter, Alexei G. Dondua, András Kosztolányi, Anne Rutten, Barbara Helm, Brett K. Sandercock, Bruce Casler, Bruno J. Ens, Caleb S. Spiegel, Chris J. Hassell, Clemens Küpper, Clive Minton, Daniel Burgas, David B. Lank, David C. Payer, Egor Y. Loktinov, Erica Nol, Eunbi Kwon, Fletcher Smith, H. River Gates, Hana Vitnerová, Hanna Prüter, James A. Johnson, James J. H. St Clair, Jean-François Lamarre, Jennie Rausch, Jeroen Reneerkens, Jesse R. Conklin, Joana Burger, Joe Liebezeit, Joël Bêty, Jonathan T. Coleman, Jordi Figuerola, Jos C. E. W. Hooijmeijer, José A. Alves, Joseph A. M. Smith, Karel Weidinger, Kari Koivula, Ken Gosbell, Klaus-Michael Exo, Larry Niles, Laura Koloski, Laura McKinnon, Libor Praus, Marcel Klaassen, Marie-Andrée Giroux, Martin Sládeček, Megan L. Boldenow, Michael I. Goldstein, Miroslav šálek, Nathan Senner, Nelli Rönkä, Nicolas Lecomte, Olivier Gilg, Orsolya Vincze, Oscar W. Johnson, Paul A. Smith, Paul F. Woodard, Pavel S. Tomkovich, Phil F. Battley, Rebecca Bentzen, Richard B. Lanctot, Ron Porter, Sarah T. Saalfeld, Scott Freeman, Stephen C. Brown, Stephen Yezerinac, Tamás Székely, Tomás Montalvo, Theunis Piersma, Vanessa Loverti, Veli-Matti Pakanen, Wim Tijsen, Bart Kempenaers

## Abstract

The behavioural rhythms of organisms are thought to be under strong selection, influenced by the rhythmicity of the environment^1–4^. Such behavioural rhythms are well studied in isolated individuals under laboratory conditions^1,5^, but free-living individuals have to temporally synchronize their activities with those of others, including potential mates, competitors, prey and predators^6–10^. Individuals can temporally segregate their daily activities (e.g. prey avoiding predators, subordinates avoiding dominants) or synchronize their activities (e.g. group foraging, communal defence, pairs reproducing or caring for offspring)^6–9,11^. The behavioural rhythms that emerge from such social synchronization and the underlying evolutionary and ecological drivers that shape them remain poorly understood^5–7,9^. Here, we address this in the context of biparental care, a particularly sensitive phase of social synchronization^12^ where pair members potentially compromise their individual rhythms. Using data from 729 nests of 91 populations of 32 biparentally-incubating shorebird species, where parents synchronize to achieve continuous coverage of developing eggs, we report remarkable within– and between-species diversity in incubation rhythms. Between species, the median length of one parent’s incubation bout varied from 1 – 19 hours, while period length–the time in which a parent’s probability to incubate cycles once between its highest and lowest value – varied from 6 – 43 hours. The length of incubation bouts was unrelated to variables reflecting energetic demands, but species relying on crypsis (the ability to avoid detection by other animals) had longer incubation bouts than those that are readily visible or actively protect their nest against predators. Rhythms entrainable to the 24-h light-dark cycle were less prevalent at high latitudes and absent in 18 species. Our results indicate that even under similar environmental conditions and despite 24-h environmental cues, social synchronization can generate far more diverse behavioural rhythms than expected from studies of individuals in captivity^5–7,9^. The risk of predation, not the risk of starvation, may be a key factor underlying the diversity in these rhythms.

---

Incubation by both parents prevails in almost 80% of non-passerine families^13^ and is the most common form of care in shorebirds^14^. Biparental shorebirds are typically monogamous^15^, mostly lay three or four eggs in an open nest on the ground^15^, and cover their eggs almost continuously^13^. Pairs achieve this through synchronization of their activities such that one of them is responsible for the nest at a given time (i.e. an incubation bout). Alternating female and male bouts generate an incubation rhythm with a specific period length (cycle of high and low probability for a parent to incubate).

We used diverse monitoring systems (Methods & Extended Data Table 1) to collect data on incubation rhythms from 91 populations of 32 shorebird species belonging to 10 genera (Fig. 1a), extracted the length of 34,225 incubation bouts from 729 nests, and determined the period length for pairs in 584 nests (see Methods, Extended Data Fig. 1 & 2).

We found vast between– and within– species variation in incubation bout length and in period length (Fig. 1–3 & Extended Data Fig. 3). Different species, but also different pairs of the same species, adopted strikingly different incubation rhythms, even when breeding in the same area (see, e.g. incubation rhythms in Barrow, Alaska, represented by in Fig. 1b-c; incubation rhythms for each nest are in Supplementary Actograms^16^). Whereas in some pairs parents exchanged incubation duties about 20 times a day (Fig. 2a); e.g. *Charadrius semipalmatus*, Fig. 1b), in others a single parent regularly incubated for 24 hours (Fig. 2a; e.g. *Limnodromus scolopaceus*, Fig. 1b), with exceptional bouts of up to 50 hours (Supplementary Actograms^16^). Similarly, whereas incubation rhythms of pairs in 22% of nests followed a strict 24-h period (Fig. 2b; e.g. *Tringa flavipes*, Fig. 1b), the rhythms of others dramatically deviated from a 24-h period (Fig. 2b) resulting in ultradian (<20-h in 12% of nests; e.g. *Numenius phaeopus*; Fig. 1b), free-running like (e.g. *Calidris alpina*; Fig. 1b) and infradian rhythms (>28-h in 8% of nests), some with period lengths up to 48-h (e.g. *Limnodromus scolopaceus*; Fig. 1b). This variation in period length partly related to the variation in bout length (Fig. 3): in the suborder Scolopaci period length correlated positively with median bout length, but in the suborder Charadrii species with 24-h periods had various bout lengths, and species with similar bout lengths had different period lengths.

**Figure 1.**
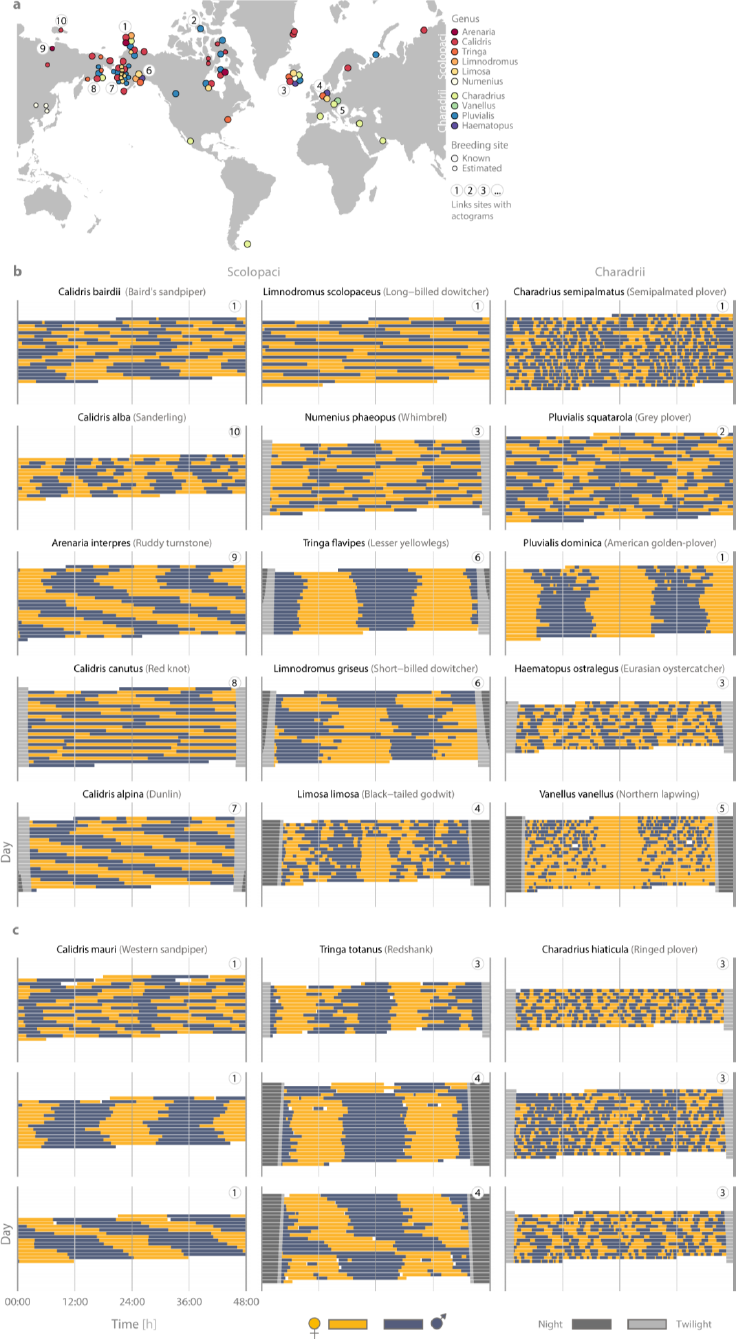
Map of studied breeding sites and the diversity of shorebird incubation rhythms. **a**, Map of breeding sites with data on incubation rhythms. The colour of the dots indicates the genus (data from multiple species per genus may be available), the size of the dots refers to data quality (◯, large: exact breeding site known; ○, small: breeding site estimated, see Methods). For nearby or overlapping locations, the dots are scattered to increase visibility. Contours of the map made with Natural Earth, http://www.naturalearthdata.com. **b-c**, Illustrations of between-species diversity (**b**) and within-species diversity (**c**; note that the three rhythms for *Calidris mauri* and *Charadrius hiaticula* come from the same breeding location.). Each actogram depicts the bouts of female (yellow; 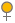) and male (blue-grey; 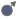) incubation at a single nest over a 24-h period, plotted twice, such that each row represents two consecutive days. If present, twilight is indicated by light grey bars (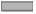) and corresponds to the time when the sun is between 6° and 0° below the horizon, night is indicated by dark grey bars (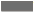) and corresponds to the time when the sun is < 6° below the horizon. Twilight and night are omitted in the centre of the actogram (24:00) to make the incubation rhythm visible. The circled numbers (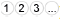) indicate the breeding site of each pair and correspond to the numbers on the map **a**.

**Figure 2.**
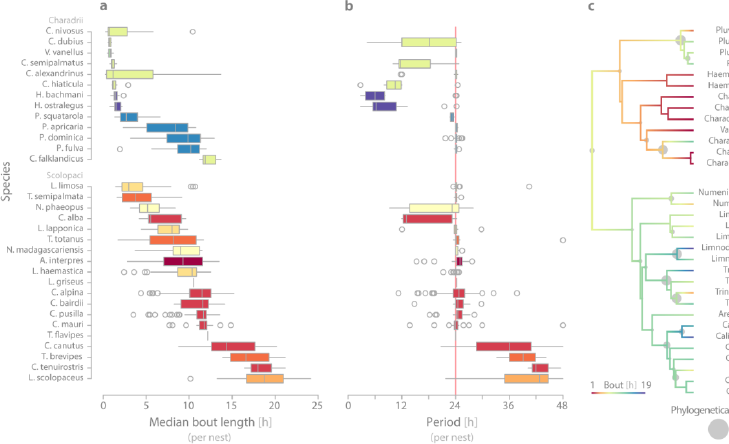
Between–and within-species variation in incubation rhythms and their estimated evolution. **a-b**, Box plots are ordered by species (within suborder) from the shortest to the longest median bout length, and depict the genus (colour as in Fig. 1a), median (vertical line inside the box), the 25^th^ and 75^th^ percentiles (box), 25^th^ percentiles minus 1.5 times interquartile range and 75^th^ percentile plus 1.5 times interquartile range or minimum/maximum value, whichever is smaller (bars), and the outliers (circles). *N*_median bout length_ = 729 and *N*_period_ = 584 nests. **b,** The red vertical line indicates a 24-h period. **c,** Observed and reconstructed incubation bout and period length visualised (by colour) on the phylogenetic tree^29^ using species’ medians (based on population medians) and one of 100 sampled trees (see Methods). The grey circles represent phylogenetically independent contrasts^30^ and hence emphasize the differences at each tree node.

**Figure 3.**
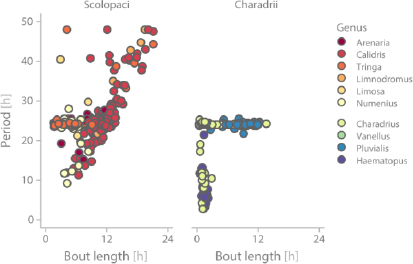
Relationship between bout and period length. Each dot represents a single nest (*N* = 584 nests), colours depict the genus. In the suborder Scolopaci the median bout length and period length correlate positively (*r_Spearman_* = 0.56, *N* = 424 nests); in the suborder Charadrii periods longer than ~24h are absent, and there is no simple relationship between bout and period length (*N* = 160 nests). For species-specific relationships see Extended Data Fig. 3.

Despite substantial within-species variation, we found a strong evolutionary signal for both bout and period length with a coefficient of phylogenetic signal λ close to 1 (Extended Data Table 2). This is consistent with the notion that biological rhythms are largely genetically determined and conserved among related species^8–10^. However, the phylogenetic effect seems unevenly distributed over taxonomic level. Suborder explained 33% of the phenotypic variance in both bout and period length, with the Scolopaci having longer incubation bouts and periods than the Charadrii (Extended Data Table 3; Fig. 2 & 3). Species explained 41% of the phenotypic variation in bout length and 46% in period length, but genus explained little (<1% in both; Extended Data Table 3), suggesting that despite a strong phylogenetic signal, these traits can rapidly diverge (Fig. 2c).

**Figure 4.**
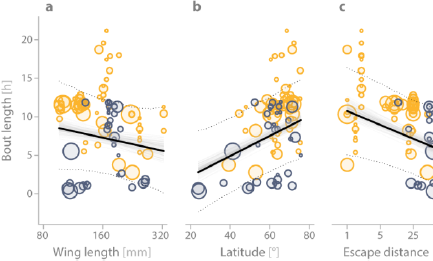
Predictors of variation in incubation rhythms. **a-c**, Relationships between bout length and body size, measured as female wing length (**a**), breeding latitude (**b**) and anti-predation strategy, quantified as escape distance (**c**) for *N* = 729 nest from 91 populations belonging to 32 species. **d,** The relationship between the proportion of nests with a period length that is entrainable by the 24-h light-dark cycle (i.e. period lengths: 3, 6, 12, 24, or 48h) and breeding latitude (*N* = 584 nests from 88 populations belonging to 30 species). **e,** The distribution of period length over latitude. The period was standardized to 24h so that all 24-h harmonics are depicted as 24h (red line) and respective deviations from each harmonic as deviations from 24h (e.g. period of 12.5h is depicted as 25h). **a-e**, Each circle represents the population median; circle size indicates the number of nests. **a-c**, The solid lines depict the model-predicted relationships, the dotted lines the 95% credible intervals based on the joint posterior distribution of 100 separate MCMCglmm runs each with one of the 100 phylogenetic trees with ~1,100 independent samples per tree. The grey areas depict the predicted relationships for each of 100 runs (i.e. the full range of regression line estimates across 100 models) and illustrate the uncertainty due to the phylogenetic tree. The predicted relationships stem from a Gaussian phylogenetic mixed-effect models, where the effects of other predictors were kept constant (**a-c**, Extended Data Table 4), or from a binomial phylogenetic mixed-effect model (**d**, Extended Data Table 4).

Two ecological factors may explain the observed variation in bout length. First, the ‘energetic demands hypothesis’ stipulates that the length of an incubation bout depends on a bird’s energetic state^13,17^. This predicts that (1) large species will have longer incubation bouts than smaller species, because they radiate less body heat per unit of mass and (2) incubation bouts will shorten with increasing breeding latitude, because – everything else being equal – energy stores will deplete faster in colder environments (Extended Data Fig. 4a-b shows latitudinal cline in summer temperatures). However, bout length was unrelated to body size (Fig. 4a) and correlated positively (instead of negatively) with latitude (Fig. 4b). These correlational results across populations and species support recent experimental findings within species^18^ and suggest that in biparentally-incubating shorebirds energetic demands are not an important ecological driver underlying variation in bout length.

An alternative explanation for variation in the length of incubation bouts relates to anti-predation strategies. Those species that rely primarily on parental crypsis (Extended data Fig. 5a) benefit from reduced activity near the nest, because such activity can reveal the nest’s location to potential predators^19,20^. Thus, in these species, selection will favour fewer change-overs at the nest, and hence longer incubation bouts. In contrast, species that are clearly visible when sitting on the nest or that rely on active anti-predation behaviour (Extended Data Fig. 5b), including having a partner on the watch for predators, leaving the nest long before the predator is nearby and mobbing the predator^15^, obtain no advantage from minimizing activity. For these species, bout length can shorten, which may be advantageous for other reasons (e.g. reduced need to store fat). We quantified anti-predation strategy as the distance at which the incubating parent left the nest when approached by a human (escape distance), because cryptic species stay on the nest longer (often until nearly stepped upon)^15^. Despite the large geographical distribution of the studied species, with related variability in the suite of predators and predation pressure^21^, and even when controlling for phylogeny (which captures much of the variation in anti-predation strategy, Extended Data Fig. 6), escape distance negatively correlated with the length of incubation bouts (Fig. 4c). This result suggests that bout length co-evolved with the anti-predation strategy.

Under natural conditions, most organisms exhibit 24-h rhythmicity, but during summer, when most shorebirds breed, the 24-h variation in light decreases with latitude leading to continuous polar daylight^22^ (Extended Data Fig. 4c-d). Such reduced variation in 24-h light intensity may cause a loss of 24-h rhythmicity^23–25^. As a consequence, circadian behavioural rhythms should exhibit a latitudinal cline^22^. As predicted, incubation rhythms with periods that do not follow the 24-h light-dark cycle, such as ‘free running-like patterns’ (left column in Fig. 1b), occurred more often in shorebirds breeding at higher latitudes (Fig. 4d).The absolute deviations of periods from 24-h and 24-h harmonics also increased with latitude (Fig. 4e; Extended Data Table 4). Although this supports the existence of a latitudinal cline in socially emerged behavioural rhythms^22^, we found a substantial number of rhythms that defy the 24-h day even at low and mid latitudes (Fig 4d-e).

Many shorebirds predominantly use tidal habitats, at least away from their breeding ground^15^. To anticipate tidal foraging opportunities, these species may have activity patterns with a period length resembling the tidal period. As changing to a different rhythm is costly^26^, these tidal activity patterns might carry over to incubation. Although half of our species are tidal away from their breeding grounds, and some forage in tidal areas also during breeding (~12% of populations), in only 5% of nests did pairs display a period length that can be entrained by the tide. Moreover, tidal species had similar (not longer) periods than non-tidal ones (Extended Data Table 4). Hence, unlike the 24-h light-dark cycle, tidal life-history seems to play at best a negligible role in determining incubation rhythms.

Three main questions arise from our results. First, is variation in incubation bout length in cryptic species related to the actual predation pressure? This can be tested by comparing bout length between populations of a particular species that are exposed to different predator densities, or between years that differ in predation pressure. Second, it remains unclear how the diverse social rhythms emerge. Are these rhythms a consequence of behavioural flexibility, or a ‘fixed’ outcome of synchronization between the circadian clocks of the two individuals involved? An experimental study on ring doves (*Streptopelia risoria*) suggests that parents may even use two timers - circadian oscillation and interval timing - to determine when to incubate^27^. Parents rapidly adjusted their schedules to phase-shifted photoperiods and their incubation rhythm ‘free-ran’ in constant dim illumination (implying a circadian mechanism), whereas an experimental delay in the onset of an incubation bout did not change the length of the bout because the incubating parent refused to leave the nest until its incubation bout reached the ‘typical’ duration (implying interval timing). Third, what – if any – are the fitness consequences for the parents of having a certain incubation rhythm? For example, the costs of having a particular incubation rhythm may be unevenly distributed between the two parents (e.g. because one parent is on incubation duty when food is more readily available, or because one parent ‘enforces’ its own rhythm at a cost to the other parent).

In conclusion, our results reveal that under natural conditions social synchronization can generate much more diverse rhythms than expected from previous work^**5-7,9,28**^, and that these rhythms often defy the assumptions of entrainment to the 24-h day-night cycle. Not risk of starvation, but risk of predation seems to play a key role in determining some of the variation in incubation rhythms. We describe this diversity in the context of biparental incubation, but diverse behavioural rhythms may also arise in many other social settings (e.g. in the context of mating interactions^25^, vigilance behaviour during group foraging). Essentially, the reported diversity suggests that the expectation that individuals within a pair (or group) should optimize their behavioural rhythms relative to the 24-h day may be too simplistic, encouraging further study of the evolutionary ecology of plasticity in circadian clocks.

**Supplementary Information** is freely available at open science framework https://osf.io/wxufm/^16^

## Acknowledgements

We thank all that made the data collection possible. We are grateful to W. Schwartz, E. Schlicht, W. Forstmeier, M. Baldwin, H. Fried Petersen, D. Starr-Glass, and B. Bulla for comments on the manuscript and to F. Korner-Nievergelt, J. D. Hadfield, L. Z. Garamszegi, S. Nakagawa, T. Roth, N. Dochtermann, Y. Araya, E. Miller and H. Schielzeth for advice on data analysis. Data collection was supported by various institutions and people listed in the Supplementary Data 1: https://osf.io/sq8gk^16^. The study was supported by the Max Planck Society (to B.K.). M.B. is a PhD student in the International Max Planck Research School for Organismal Biology.

## Author Contributions

M.B. and B.K. conceived the study. All authors except B.H. collected the primary data. MB coordinated the study and managed the data. MB and M.V. developed the methods to extract incubation. M.B. extracted bout lengths and with help from A.R. and M.V. created actograms. M.B. with help from M.V. analysed the data. M.B. prepared the Supporting Information. M.B. and B.K. wrote the paper with input from the other authors. Except for the first, second, and last author, the authors are listed alphabetically by their first name.

## Author Information

All information, primary and extracted data, computer code and software necessary to replicate our results, as well as the Supplementary Actograms are open access and archived at Open Science Framework https://osf.io/wxufm/^16^. The authors declare no competing financial interests. Correspondence and requests for materials should be addressed to M.B. and B.K.

## METHODS

**Recording incubation.** Incubation data were obtained between 1994 and 2015, for as many shorebird species (*N* = 32) and populations (*N* = 91) as possible, using six methods (for specifications of the equipment see Extended Data Table 1). (1) In 261 nests, a radio frequency identification reader (‘RFID’) registered presence of tagged parents at the nest. The passive-integrated tag was either embedded in a plastic flag^31,32^, with which the parents were banded, or glued to the tail feathers^33^. In 200 nests the RFID was combined with a temperature probe placed between the eggs. The temperature recordings allowed us to identify whether a bird was incubating even in the absence of RFID readings; an abrupt change in temperature demarcated the start or end of incubation^31^. (2) For 396 nests, light loggers were mounted to the plastic flag or a band that was attached to the bird’s leg^34,35^. The logger recorded maximum light intensity (absolute or relative) for a fixed sampling interval (2-10 min). An abrupt change in light intensity (as opposed to a gradual change caused, e.g. by civil twilight) followed by a period of low or high light intensity demarcated the start or end of the incubation period (Extended Data Fig. 2). (3) For nine nests a GPS tag, mounted on the back of the bird, recorded the position of the bird^36^. The precision of the position depends on cloud cover and sampling interval^36^. Hence, to account for the imprecision in GPS positions, we assumed incubation whenever the bird was within 25 m of the nest (Extended Data Fig. 2b). (4) At three nests automated receivers recorded signal strength of a radio-tag attached to the rump of a bird; whenever a bird incubated, the strength of the signal remained constant^24^ (Supplementary Actograms p. 257-9^16^). (5) At 53 nests video cameras and (6) for 8 nests continuous observations were used to identify the incubating parents; parent identification was based on plumage, colour rings or radio-tag. In one of the populations, three different methods were used, in seven populations representing seven species two methods. In one nest, two methods were used simultaneously (Extended Data Fig. 2b).

### Extraction of incubation bouts

An incubation bout was defined as the total time allocated to a single parent (i.e. the time between the arrival of a parent at and its departure from the nest followed by incubation of its partner). Bout lengths were only extracted if at least 24h of continuous recording was available for a nest; in such cases, all bout lengths were extracted. For each nest, we transformed the incubation records to local time as (UTC time + 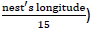). Incubation bouts from RFIDs, videos and continuous observations were mostly extracted by an R-script and the results verified by visualizing the extracted and the raw data^16,31,37,38^; otherwise, MB extracted the bouts manually from plots of raw data^39,40^ (plots of raw data and extracted bouts for all nests are in the Supplementary Actograms^16^; the actograms were generated by ‘ggplot’ and ‘xyplot’ functions from the ‘ggplot2’ and ‘lattice’ R package^41–43^). Whenever the start or end of a bout was unclear, we classified these bouts as uncertain (see next paragraph for treatment of unsure bouts). In case of light logger data, the light recordings before and after the breeding period, when the birds were definitely not incubating, helped to distinguish incubation from non-incubation. Whenever an individual tagged with a light logger nested in an environment where the sun was more than 6° below the horizon for part of a day (i.e. night), we assumed an incubation bout when the individual started incubating before the night started and ended incubating after the night ended. When different individuals incubated at the beginning vs. at the end of the night, we either did not quantify these bouts or we indicated the possible time of exchange (based on trend in previous exchanges), but classified these bouts as uncertain (see Supplementary Actograms^16^). In total, we extracted 34,225 incubation bouts.

The proportion of uncertain bouts within nests had a distribution skewed towards zero (median = 0%, range: 0-100%, *N* = 729 nests), and so did the median proportion of uncertain bouts within populations (median = 2%, range: 0-74%, *N* = 91 populations). Excluding the uncertain bouts did not change our estimates of median bout length (Pearson’s correlation coefficient for median bout length based on all bouts and without uncertain bouts: *r* = 0.96, *N* = 335 nests with both certain and uncertain bouts). Hence, in further analyses all bouts were used to estimate median bout length.

Note that in some species sexes consistently differ in bout length (Figure 1b, e.g. *Vanellus vanellus*). As these differences are small compared to the between-species differences and because in 27 nests (of 8 species) the sex of the parents was unknown, we here use median bout length independent of sex.

### Extraction of period length

The method used for extracting the period length of incubation rhythm for each nest is described in the Extended Data Fig. 1.

### Extraction of entrainable periods

We classified 24-h periods and periods with 24-h harmonics (i.e. 3, 6, 12, 48h) as strictly entrainable by 24-h light fluctuations (*N* = 142 nests out of 584). Including also nearest adjacent periods (±0.25h) increased the number of nests with entrainable periods (*N* = 277), but results of statistical analyses remained quantitatively similar. We consider periods and harmonics of 12.42h (i.e. 3.1, 6.21, 12.42, 24.84h) as strictly entrainable by tide. However, because the periods in our data were extracted in 0.25-h intervals (Extended Data Fig. 1), we classified periods of 3, 6.25, 12.5, 24.75h (i.e. those closest to the strict tide harmonics) as entrainable by tide (*N* = 32 nests out of 584). Including also the second nearest periods (i.e. 3.25, 6, 12.25, 25) increased the number of nests entrainable by tide to *N* = 55.

### Population or species life-history traits

For 643 nests, the exact breeding location was known (nests or individuals were monitored at the breeding ground). For the remaining 86 nests (from 27 populations representing 8 species, where individuals were tagged with light loggers on the wintering ground), the breeding location was roughly estimated from the recorded 24-h variation in daylight, estimated migration tracks, and the species’ known breeding range^44–51^. One exact breeding location was in the Southern Hemisphere, so we used absolute latitude in analyses. Analyses without populations with estimated breeding-location or without the Southern Hemisphere population generated quantitatively similar estimates as the analyses on full data.

For each population, body size was defined as mean female wing length^52^, either for individuals measured at the breeding area or at the wintering area. In case no individuals were measured, we used the mean value from the literature (see open access data for specific values and references^53^).

Anti-predation strategy was assessed by estimating escape distance of the incubating bird when a human approached the nest, because species that are cryptic typically stay on the nest much longer than non-cryptic species, sometimes until nearly stepped upon^48,54^. Escape distance was obtained for all species. Forty-four authors of this paper estimated the distance (in m) for one or more species based on their own data or experience. For 10 species, we also obtained estimates from the literature^48^. We then used the median ‘estimated escape distance’ for each species. In addition, for 13 species we obtained ‘true escape distance’. Here, the researcher approached a nest (of known position) and either estimated his distance to the nest or marked his position with GPS when the incubating individual left the nest. For each GPS position, we calculated the Euclidian distance from the nest. In this way we obtained multiple observations per nest and species, and we used the median value per species (weighted by the number of estimates per nest) as the ‘true escape distance’. The species’ median ‘estimated escape distance’ was a good predictor of the ‘true escape distance’ (Pearson’s correlation coefficient: *r* = 0.89, *N* = 13 species). For analysis, we defined the escape distance of a species as the median of all available estimates.

For each species, we determined whether it predominantly uses a tidal environment outside its breeding ground, i.e. has tidal vs. non-tidal life history (based on^48,50,51^). For each population with exact breeding location, we scored whether tidal foraging habitats were used by breeding birds for foraging (for three populations this information was unknown)^53^. For all populations with estimated breeding location we assumed, based on the estimated location and known behaviour at the breeding grounds, no use of tidal habitat.

### Statistical analyses

Unless specified otherwise, all analyses were performed on the nest level using median bout length and extracted period length.

We used phylogenetically informed comparative analyses to assess how evolutionary history constrains the incubation rhythms (estimated by Pagel’s λ coefficient of phylogenetic signal^55,56^) and to control for potential non-independence among species due to common ancestry. This method explicitly models how the covariance between species declines as they become more distantly related^55,57,58^. We used the Hackett^59^ backbone phylogenetic trees available at http://birdtree.org^60^, which included all but one species (*Charadrius nivosus*) from our dataset. Following a subsequent taxonomic split^61^, we added *Charadrius nivosus* to these trees as a sister taxon of *Charadrius alexandrinus*. Phylogenetic uncertainty was accounted for by fitting each model with 100 phylogenetic trees randomly sampled from 10,000 phylogenies at http://birdtree.org^60^.

The analyses were performed with Bayesian phylogenetic mixed-effect models (Fig. 4 and Extended Data Table 2 and 4) and the models were run with the ‘MCMCglmm’ function from the R package ‘MCMCglmm’^62^. In all models, we also accounted for multiple sampling within species and breeding site (included as random effects). In models with a Gaussian response variable, an inverse-gamma prior with shape and scale equal to 0.001 was used for the residual variance (i.e. variance set to one and the degree of belief parameter to 0.002). In models with binary response variables, the residual variance was fixed to one. For all other variance components the parameter-expanded priors were used to give scaled F-distributions with numerator and denominator degrees of freedom set to one and a scale parameter of 1,000. Model outcomes were insensitive to prior parameterization. The MCMC chains ran for 2,753,000 iterations with a burn-in of 3,000 and a thinning interval of 2,500. Each model generated ~1,100 independent samples of model parameters (Extended Data Table 2 and 4). Independence of samples in the Markov chain was assessed by tests for autocorrelation between samples and by using graphic diagnostics.

First, we used MCMCglmm to estimate Pagel’s λ (phylogenetic signal) for bout and period length (Gaussian), and to show that our estimates of these two incubation variables were independent of how often the incubation behaviour was sampled (‘sampling’ in min, ln-transformed; Extended Data Table 2). Hence, in subsequent models, sampling was not included.

Then, we used MCMCglmm to model variation in bout length and period length. Bout length was modelled as a continuous response variable and latitude (°, absolute), female wing length (mm, ln-transformed) and approach distance (m, ln-transformed) as continuous predictors. Predictors had low collinearity (at nest, population and species level; all Pearson or Spearman correlation coefficients |*r*| < 0.28). To test for potential entrainment to 24-h, period length was modelled as a binary response variable (1 = rhythms with period of 3, 6, 12, 24, or 48 h; 0 = rhythms with other periods) and latitude as a continuous predictor. To test how circadian period varies with latitude or life history, period was transformed to deviations from 24-h and 24-h harmonics and scaled by the time span between the closest harmonic and the closest midpoint between two harmonics. For example, a 42h period deviates by- 6h from 48h (the closest 24-h harmonic) and hence -6h was divided by 12h (the time between 36h – the midpoint of two harmonics - and 48h -the closest harmonic). This way the deviations spanned from -1 to 1 with 0 representing 24-h and its harmonics. The absolute deviations were then modelled as a continuous response variable and latitude as continuous predictor. The deviations were also modelled as a continuous response and species life history (tidal or not) as categorical predictor.

In all models the continuous predictor variables were centred and standardized to a mean of zero and a standard deviation of one.

We report model estimates for fixed and random effects, as well as for Pagel’s λ, by the modes and the uncertainty of the estimates by the highest posterior density intervals (referred to as 95% CI) from the joint posterior distributions of all samples from the 100 separate runs, each with one of the 100 separate phylogenetic trees from http://birdtree.org.

To help interpret the investigated relationships we assessed whether incubation rhythms evolved within diverged groups of species by plotting the evolutionary tree of the incubation rhythm variables (Fig. 2c), as well as of the predictors (Extended Data Fig. 6).

The source of phylogenetic constraint in bout and period length was investigated by estimating the proportion of phenotypic variance explained by suborder, genus and species (Extended Data Table 3). The respective mixed models were also specified with ‘MCMCglmm’^62^ using the same specifications as in the phylogenetic models. Because suborder contained only two levels, we first fitted an intercept mixed model with genus, species, and breeding site as random factors, and used it to estimate the overall phenotypic variance. We then entered suborder as a fixed factor and estimated the variance explained by suborder as the difference between the total variance from the first and the second model. To evaluate the proportion of the variance explained by species, genus and breeding site, we used the estimates from the model that included suborder.

R version 3.1.1^63^ was used for all statistical analyses.

### Code availability

All statistical analyses, figures, and the Supplementary Actograms are replicable with the open access information, software and r-code available from Open Science Framework, https://osf.io/wxufm/^16^.

### Data availability

Primary and extracted data that support the findings of this study are freely available from Open Science Framework, https://osf.io/wxufm/^16^.

## EXTENDED DATA

**Extended Data Figure 1.**
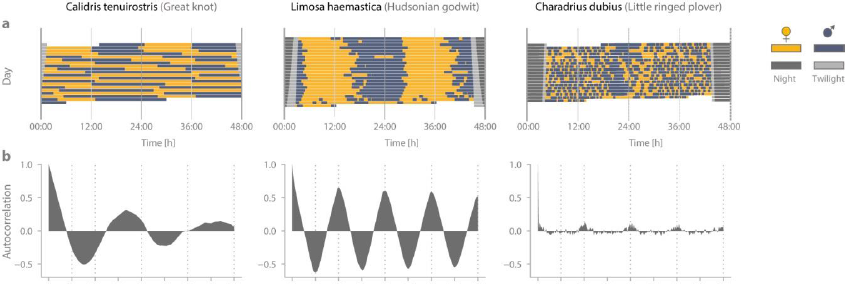
Extracting period length of incubation rhythms. **a-c,** Each column represents an example for a specific nest with long, intermediate and short incubation bouts. **a**, From the extracted bout lengths we created a time series that indicated for each nest, and every 10 min, whether a specific parent (female, if sex was known) incubated or not. Exchange gaps (no parent on the nest) had to be < 6 h to be included (for treatment of exchange gaps > 6 h see **d**, **e**). **b**, We then estimated the autocorrelation for each 10 min time-lag up to 4 days (R ‘acf’ function^63^). Positive values indicate a high probability that the female was incubating, negative values indicate that it was more likely that the male was incubating. We used only nests that had enough data to estimate the autocorrelation pattern (*N* = 584 nests from 88 populations of 30 species). The visualized autocorrelation time series never resembled white or random noise indicative of an arrhythmic incubation pattern. To determine the period (i.e. cycle of high and low probability for a parent to incubate) that dominated the incubation rhythm, we fitted to the autocorrelation estimates a series of periodic logistic regressions. In each regression, the time lag (in hours) transformed to radians was represented by a sine and cosine function 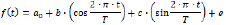, where *f*(*t*) is the autocorrelation at time-lag *t*, *a*_0_ is the intercept, *b* is the slope for sine and *c* the slope for cosine, *T* represents the length of the fitted period (in hours), and *e* is an error term. We allowed the period length to vary from 0.5 h to 48 h (in 15 min intervals, giving 191 regressions). **c**, By comparing the Akaike’s Information Criterion^64^ (AIC) of all regressions, we estimated, for each nest, the length of the dominant period in the actual incubation data (best fit). Regressions with ΔAIC (AIC_model_-AIC_min_) close to 0 are considered as having strong empirical support, while models with ΔAIC values ranging from 4-7 have less support^64^. In 73% of all nests, we determined a single best model with ΔAIC <= 3 (**c**, middle ΔAIC graph), in 20% of nests two best models emerged and in 6% of nests 3 or 4 models had ΔAIC <= 3 (**c**, left and right ΔAIC graphs). However, in all but three nests, the models with the second, third, etc. best ΔAIC were those with period lengths closest to the period length of the best model (**c**, left and right ΔAIC graphs). This suggests that multiple periodicities are uncommon. **d-e**, The extraction of the period length (described in **a-c**) requires continuous datasets, but some nests had long (>6 h) gaps between two consecutive incubation bouts, for example because of equipment failure or because of unusual parental behaviour. In such cases, we excluded the data from the end of the last bout until the same time the following day, if data were then available again (**d**), or we excluded the entire day (**e**).

**Extended Data Figure 2.**
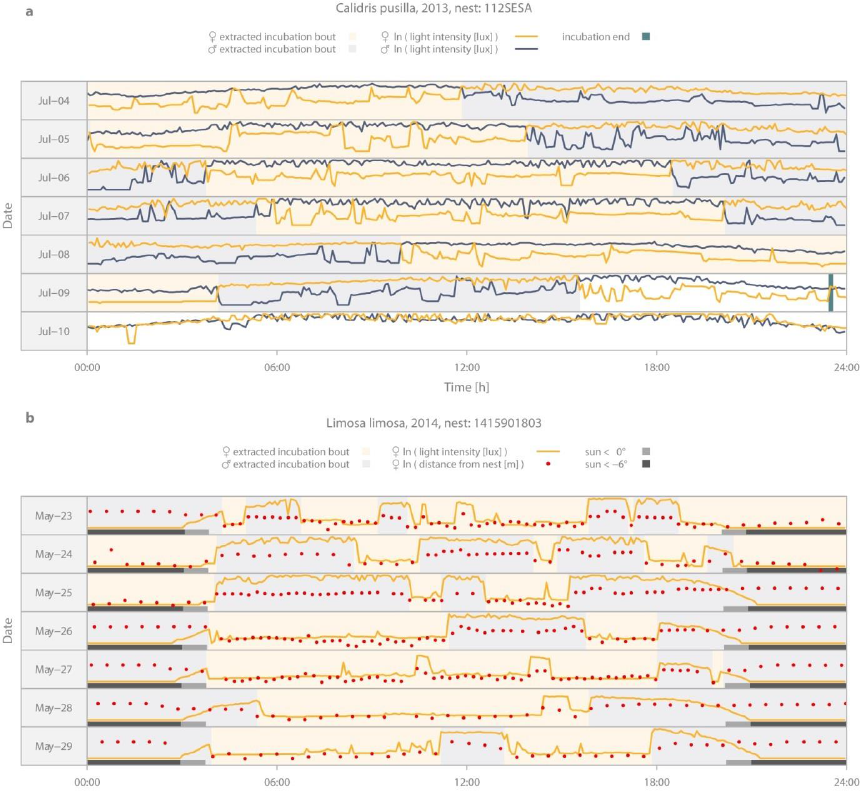
Extracting incubation bouts from light logger data. **a,** An example of a nest with a light intensity signal from both parents (yellow line 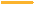 and blue-grey line 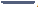). The incubation bouts for a given parent reflect periods dominated by lower light values compared to those of the partner. Note the sharp drop in the light levels at the beginning of each incubation bout and the sharp increase in the light levels at the end. Change-overs between partners occur when the light signal lines cross. Such pronounced changes in light intensity detected by the logger were used to assign incubation even when only a single parent was tagged. Note that after the chicks hatch and leave the nest (July 9, vertical bar), the light intensity signal from both parents remains similar. **b,** An example of a nest where one incubating parent was simultaneously equipped with a light logger and with a GPS tag. The yellow line (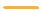) indicates light levels, red dots (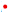) indicate the distance of the bird to the nest. As expected, low light levels co-occur with close proximity to the nest, and hence reflect periods of incubation. Although light levels decrease during twilight (light grey horizontal bar; 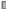), the recordings were still sensitive enough to reflect periods of incubation, i.e. the light signal matches the distance (e.g. May-25: female incubated during dawn, but was off the nest during dusk). **a-b**, Rectangles in the background indicate extracted female (light yellow polygon,) and male (light blue-grey polygon,) incubation bouts.

**Extended Data Figure 3.**
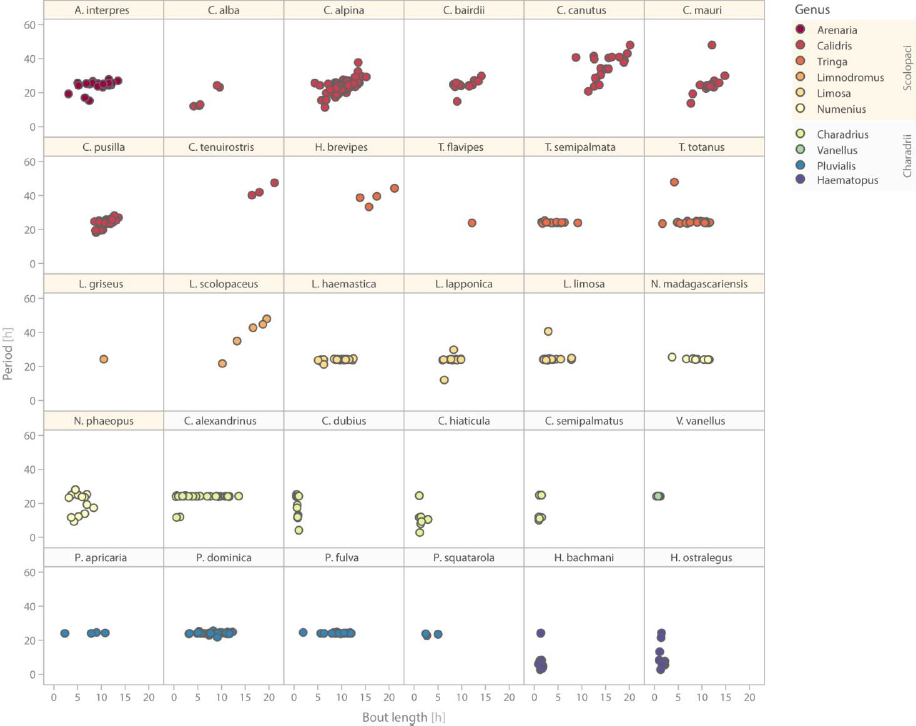
Relationship between bout and period length for 30 shorebird species. Each dot represents one nest (*N* = 584 nests), colours indicate the genus.

**Extended Data Figure 4.**
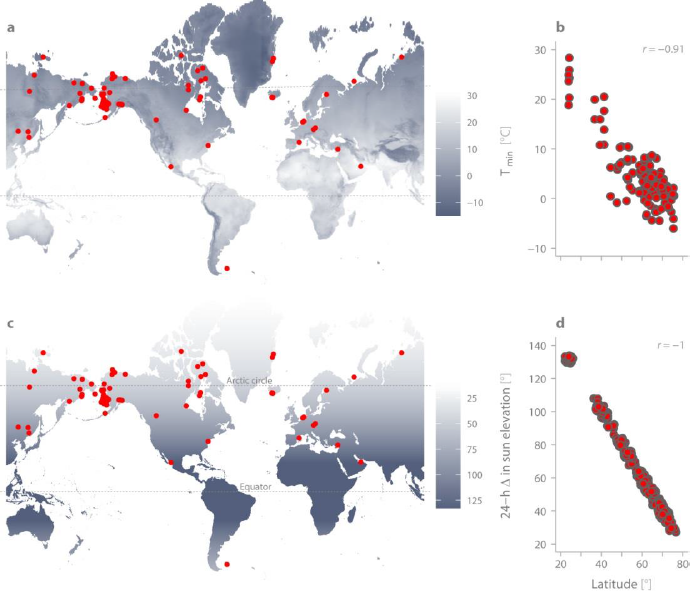
Ecological correlates of latitude. **a**, Variation in minimum temperature across the globe represented by mean minimum June temperature for the Northern Hemisphere and mean minimum December temperature for the Southern Hemisphere. **b**, Correlation between absolute latitude and the mean minimum temperature based on the month represented by mid-day of incubation data for each nest (*N* = 729). For maximum temperature the correlation was similar (r =-0.91, *N* = 729 nests). **c**, Daily variation in sun elevation (i.e. in light conditions) represented as the difference between the noon and midnight sun-elevation for the summer solstice in the Northern Hemisphere and the winter solstice in the Southern Hemisphere. **d**, Correlation between absolute latitude and daily variation in sun elevation for mid-day of incubation data for each nest (*N* = 729 nests). The points are jittered, as else they form a straight line. **a, c**, Red points indicate the breeding sites (*N* = 91). **a-b**, The minimum and maximum monthly temperature data were obtained from www.worldclim.org using the ‘raster’ R-package^65^.**c-d**, Sun-elevation was obtained by the ‘solarpos’ function from the ‘maptools’ R-package^66^.

**Extended Data Figure 5.**
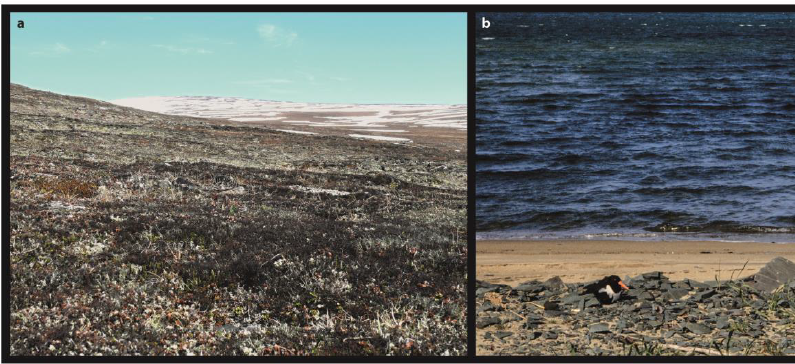
Between-species variation in parental crypsis during incubation. **a-b**, Shorebirds vary in how visible they are on the nest while incubating. The nearly invisible Great knot (*Calidris tenuirostris*; **a**; central and facing right) sits tight on the nest when approached by a human until nearly stepped upon. In contrast, the conspicuous Eurasian oystercatcher (*Haematopus ostralegus*; **b**) is visible on the nest from afar and when approached by a human leaves the nest about 100 m in advance (Credits: **a**, M. šálek; **b**, Jan van de Kam).

**Extended Data Figure 6.**
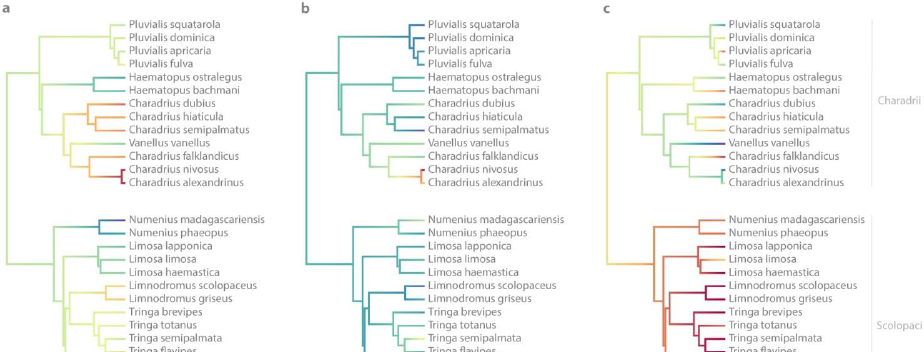
Phylogenetic relationships for predictors. **a**, Body size, estimated as female wing length. **b**, Latitude (absolute), **c**, Escape distance. **a-c**, We visualised the evolution of these traits^29,67^ using species’ medians (**a-b**; based on population medians), species’ estimates of escape distance (**c**) and one of the 100 sampled trees (see Methods).

**Extended Data Table 1.**
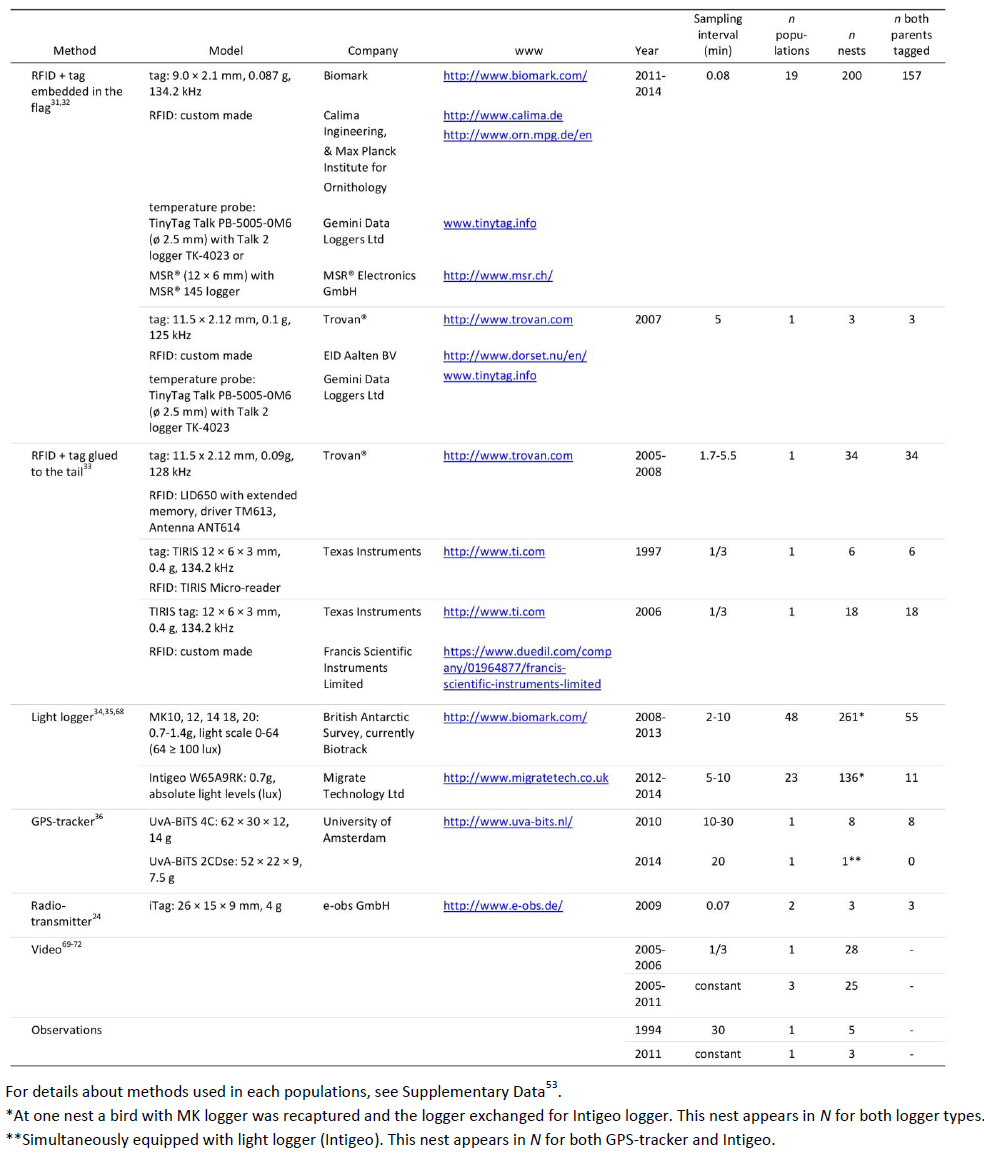
**Incubation monitoring methods and systems.**

**Extended Data Table 2.**
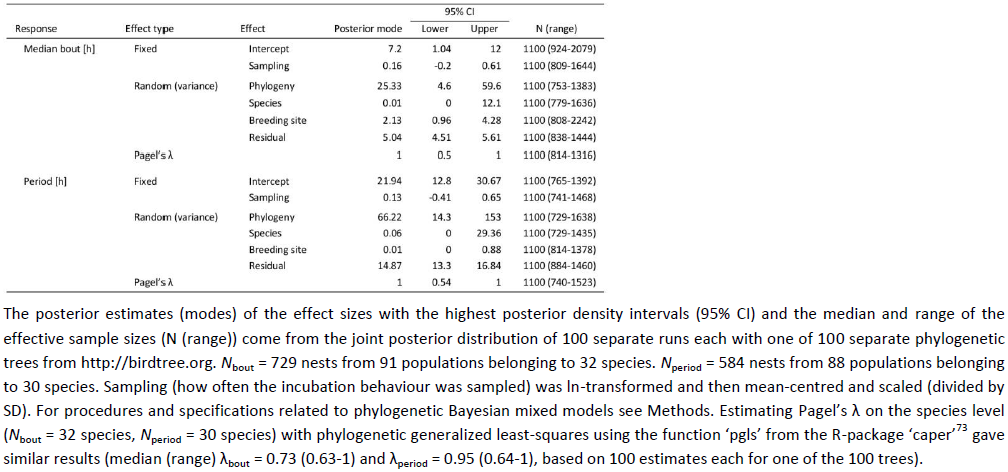
**Effects of phylogeny and sampling on bout length and period length.**

**Extended Data Table 3.**
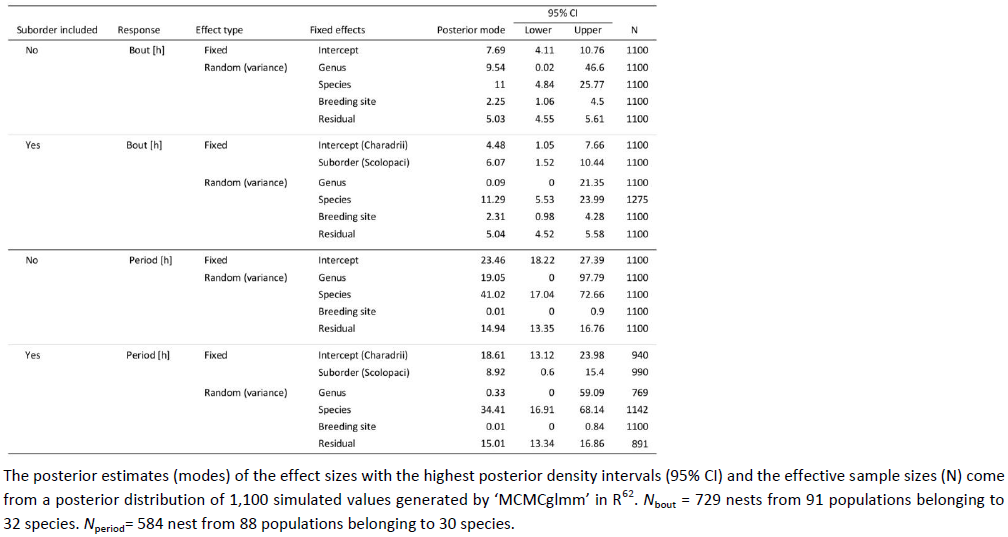
**Source of phylogenetic signal**

**Extended Data Table 4.**
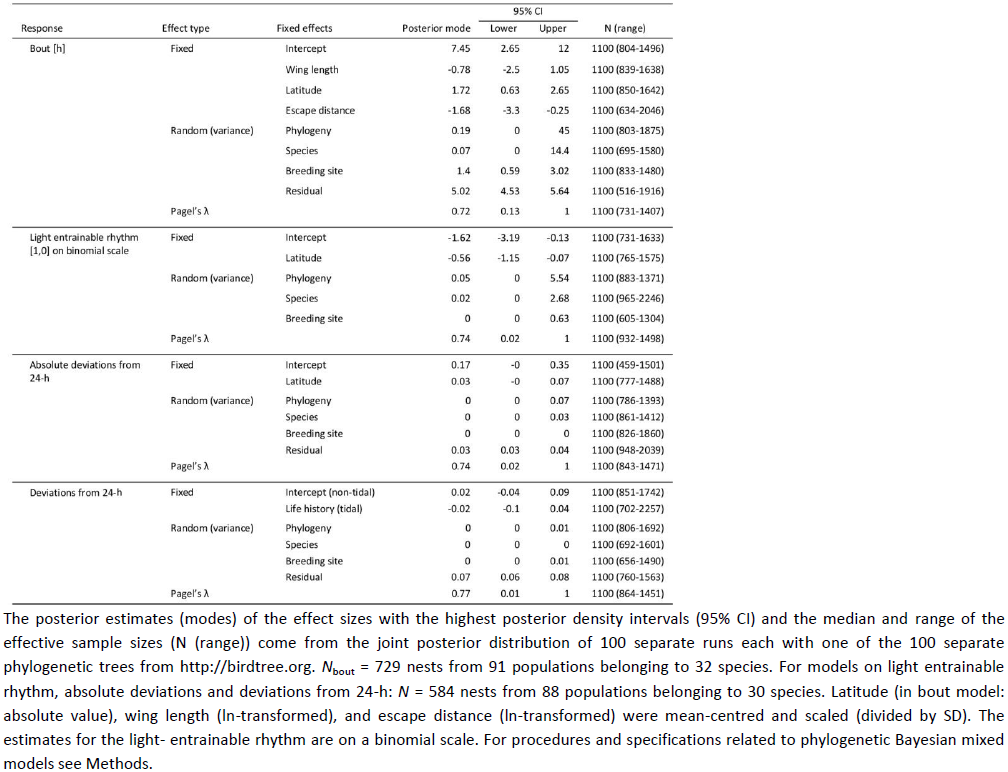
**Effect of latitude, body size, escape distance and life history on biparental incubation rhythms in shorebirds**

